# A *ZIP1* Separation-of-Function Allele Reveals that Meiotic Centromere Pairing Drives Meiotic Segregation of Achiasmate Chromosomes in Budding Yeast

**DOI:** 10.1101/265652

**Authors:** Emily L. Kurdzo, Hoa H Chuong, Dean S. Dawson

## Abstract

In meiosis I, homologous chromosomes segregate away from each other - the first of two rounds of chromosome segregation that allow the formation of haploid gametes. In prophase I, homologous partners become joined along their length by the synaptonemal complex (SC) and crossovers form between the homologs to generate links called chiasmata. The chiasmata allow the homologs to act as a single unit, called a bivalent, as the chromosomes attach to the microtubules that will ultimately pull them away from each other at anaphase I. Recent studies, in several organisms, have shown that when the SC disassembles at the end of prophase, residual SC proteins remain at the homologous centromeres providing an additional link between the homologs. In budding yeast, this centromere pairing is correlated with improved segregation of the paired partners in anaphase. However, the causal relationship of prophase centromere pairing and subsequent disjunction in anaphase has been difficult to demonstrate as has been the relationship between SC assembly and the assembly of the centromere pairing apparatus. Here, a series of in-frame deletion mutants of the SC component Zip1 were used to address these questions. The identification of separation-of-function alleles that disrupt centromere pairing, but not SC assembly, have made it possible to demonstrate that centromere pairing and SC assembly have mechanistically distinct features and that prophase centromere pairing function of Zip1 drives disjunction of the paired partners in anaphase I.

**AUTHOR SUMMARY:** The generation of gametes requires the completion of a specialized cell división called meiosis. This division is unique in that it produces cells (gametes) with half the normal number of chromosomes (such that when two gametes fuse the normal chromosome number is restored). Chromosome number is reduced in meiosis by following a single round of chromosome duplication with two rounds of segregation. In the first round, meiosis I, homologous chromosomes first pair with each other, then attach to cellular cables, called microtubules, that pull them to opposite sides of the cell. It has long been known that the homologous partners become linked to each other by genetic recombination in a way that helps them behave as a single unit when they attach to the microtubules that will ultimately pull them apart. Recently, it was shown, in budding yeast and other organisms, that homologous partners can also pair at their centromeres. Here we show that this centromere pairing also contributes to proper segregation of the partners away from each other at meiosis I, and demonstrate that one protein involved in this process is able to participate in multiple mechanisms that help homologous chromosomes to pair with each other before being segregated in meiosis I.

## INTRODUCTION

In meiosis I, homologous chromosomes segregate away from each other - the first of two rounds of segregation that allow the formation of haploid gametes. In order to segregate from one another the homologs must first become tethered together as a unit, called a bivalent. As a single bivalent, the partners can attach to microtubules such that the centromeres of the homologs will be pulled towards opposite poles of the spindle at the first meiotic division. Crossovers between the aligned homologs provide critical links, called chiasmata, which allow the homologs to form a stable bivalent (reviewed in (1)). Failures in crossing-over are associated with elevated levels of meiotic segregation errors in many organisms, including humans (reviewed in (2)). However, there are mechanisms, other than crossing-over, that can also tether partner chromosomes. Notably, studies in yeast and mouse spermatocytes have revealed that the centromeres of partner chromosomes pair in prophase of meiosis I (3-6). In budding yeast, it has been shown that this centromere pairing is correlated with the proper segregation of chromosome pairs that have failed to form chiasmata. But the formal demonstration that centromere pairing in prophase directly drives disjunction in anaphase has been difficult, because the mutations that disrupt centromere pairing also disrupt other critical meiotic processes (7, 8).

The protein Zip1 in budding yeast localizes to paired centromeres in meiotic prophase and is necessary for centromere pairing (Fig. 1A) (7-10), and similar observations have been made in Drosophila oocytes and mouse spermatocytes (3, 6, 11). Zip1 is expressed early in meiosis and first appears as dispersed punctate foci in the nucleus. Some, but not all, of these foci co-localize with centromeres, and indeed, Zip1 mediates the homology-independent pairing of centromeres at this stage of meiosis, a phenomenon called centromere-coupling (Fig. 1 A, green arrowhead) (10, 12). Zip1 later acts as a component of the synaptonemal complex (SC) - a proteinaceous structure that assembles between the axes of the homologous partners as they become aligned in meiotic prophase (Fig.1 A, blue arrowhead) (13). In budding yeast and mouse spermatocytes, when the SC disassembles in late prophase Zip1/SYCP1 remains at the paired centromeres, leaving the homologous partners only visibly joined by chiasmata and centromere pairing (Fig. 1) (3, 6-8). Most Zip1/SYCP1 appears to have left the chromosomes by the time they begin attaching to the meiotic spindles. The prophase association promoted by Zip1 is correlated with proper segregation, as *zip1* deletion mutants have no centromere pairing and also segregate achiasmate partners randomly (Fig. 1A) (7, 8).

**Figure 1.**
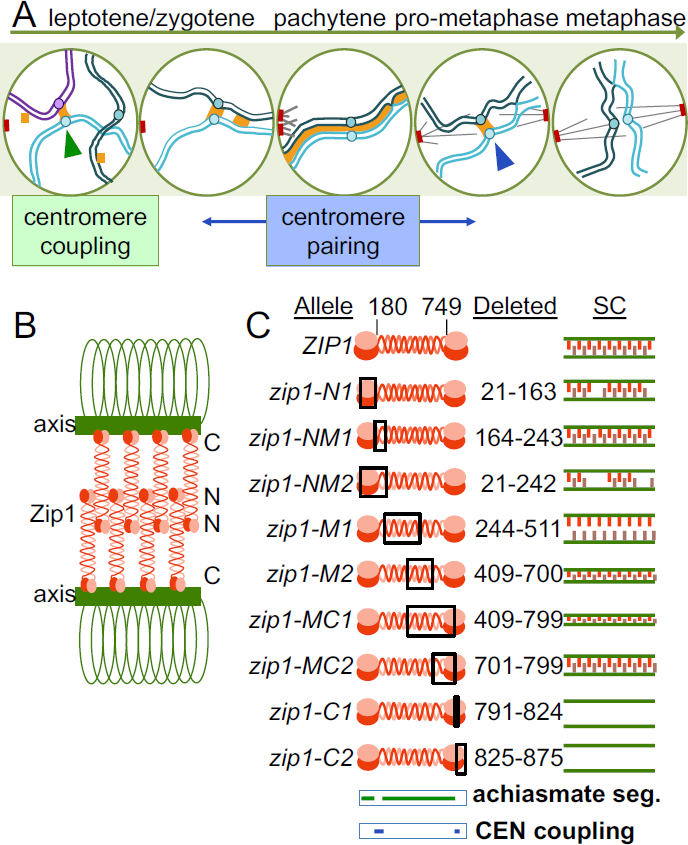
**Meiotic centromere behaviors in budding yeast. A.** In meiosis of budding yeast, Zip1 (orange) mediates centromere coupling (green arrowheads) between non-homologous partner chromosomes (light blue and purple). As the cell proceeds through later stages of meiosis, homologs pair and the mature synaptonemal complex (SC) structure zips the chromosomes together. After pachytene, the SC disassembles, except at the centromeres (blue arrowhead). **B.** The Zip1 protein is predicted to have globular domains at its ends spanning a longer coiled-coil and forms parallel dimers with N-termini in the center of the SC (denoted by N) and the C-termini along the axial elements (denoted by C). **C**. We evaluated the same nine *ZIP1* deletion mutants previously described by Tung and colleagues (Tung & Roeder, 1998). The mutations are named for their relative position along the genetic sequence N for N-terminus, M for middle region, and C for C-terminus. The approximate SC structure formed in each mutant as described by Tung and Roeder 1998 is shown.

A critical study by Tung & Roeder identified functional domains of Zip1 that contribute to SC assembly, and contributed to the current model for the structure of the SC (14). This and other studies (15) have suggested that in the SC, Zip1 is in the form of head-to-head dimers (Fig. 1 B). These dimers, in turn are thought to assemble in a ladder-like structure with the N-termini in the center of the SC and the C-termini associated with the axes of the homologous partners (Fig. 1 B). This model has been extrapolated to other organisms because the basic structure of transverse filament components, like Zip1, are believed to be conserved even though their amino acid sequences have diverged (reviewed in (16)).

Tung and Roeder 1998 used an ordered series of in-frame deletions of *ZIP1* to identify ways in which different regions of the protein contributed to SC structure and function (Fig. 1 C). This was before the discovery that Zip1 is also involved in promoting centromere coupling and centromere pairing. We have re-constructed this deletion series to evaluate the ways in which different regions of Zip1 contribute to these centromere-associated functions. This information could be used to reveal relationships in the underlying mechanisms of centromere coupling, centromere pairing and SC assembly, and identify to separation-of-function alleles that would reveal more specifically contributions made to these processes by Zip1. These approaches make clear that centromere coupling, centromere pairing, and SC assembly all require certain parts of the Zip1 protein that are not required by the others suggesting mechanistic differences in these phenomena. Second, they provide a clear demonstration that centromere pairing in prophase, distinct from other SC-related functions of Zip1, drives disjunction of achiasmate partner chromosomes in anaphase I.

## RESULTS

### The N and C terminal globular domains of Zipl are essential for centromere coupling

A series of nine in-frame deletion mutants (Fig. 1 C) were tested to determine which regions of the *ZIP1* coding sequence are essential for the homology independent centromere coupling that occurs in early meiotic prophase. Centromere coupling was assayed by monitoring the numbers of kinetochore foci (Mtw1-MYC) in chromosome spreads from prophase meiotic cells (10, 12) (Fig. 2 A). Diploid yeast have sixteen pairs of homologous chromosomes. When the centromeres of the thirty-two chromosomes are coupled they form on average sixteen Mtw1 MYC foci (Fig. 2 B, *ZIP1,* blue line). Mutants that are defective in coupling exhibit higher numbers of Mtw1-MYC foci (Fig. 2 B, *zip1Δ,* red line). The experiment was done in strains lacking *SPO11,* which encodes the endonuclease responsible for creating programmed double strand DNA (17)). This blocks meiotic progression beyond the coupling stage and prevents the homologous alignment of chromosomes (12, 18). The strains also featured GFP-tagged copies of the centromeres of chromosome I. Briefly, 256 repeats of the *lac* operon sequence was inserted adjacent to the centromere of chromosome I *(CEN1)* and the cells were engineered to express lacI-GFP, which localizes to the lacO array (19). In the centromere coupling stage, the two CEN1-GFP foci are nearly always separate because coupling is usually between non homologous partner chromosomes (Fig. 2 A) (10).

**Figure 2.**
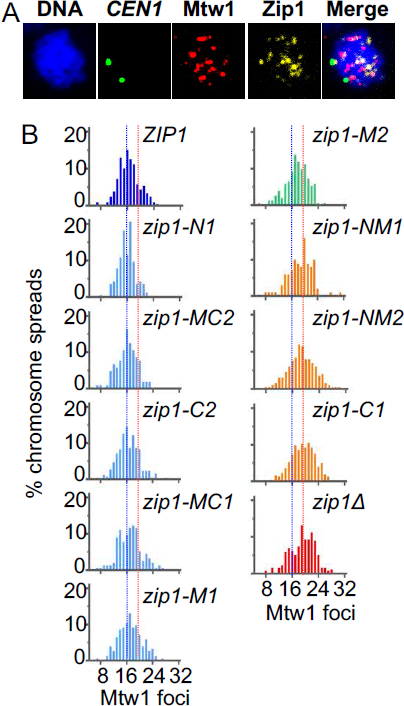
**Centromere coupling requires parts of the N and C-termini of Zip1. A.** Centromere coupling values were obtained by scoring the number of Mtw1-GFP foci in meiotic chromosome spreads. *CEN1* loci were visualized by virtue of lacI-GFP localized to a *lac* operator array next to the centromere. **B.** Coupling data. Mutants are listed according to the severity of their coupling phenotype. The thin blue and red lines indicate average Mtw1 foci values for wild-type and *ziplΔ,* respectively. The mutants were split into three groups like wild-type (light blue), intermediate (green), and like *zip1Δ* (orange). The “like wild-type” group had values indistinguishable from wild-type but were significantly different from *zip1Δ* (p<0.05); whereas the “like *zip1Δ”* group had values indistinguishable from *zip1Δ* but significantly different from wild-type (p<0.05). The *zip1-M2* mutant had an intermediate phenotype that was significantly different from both wild-type and *zip1Δ.* A complete list of averages and statistical values are presented in Table S2.

The mutants could be assigned to one of three groups based on their coupling phenotypes (Fig. 2 B and Supplemental Table 2), indistinguishable from *ZIP1* (proficient for coupling; blue histograms), indistinguishable from *zip1Δ* (loss of coupling; red and orange histograms), or intermediate (green histogram) (Fig. 2 B). The results make it possible to assign functional roles to several portions of Zip1. First, a portion of the N-terminus and adjacent coiled-coil (NM1 region, amino acids 164-242) is critical for centromere coupling. This region was shown to be largely dispensable for SC assembly and sporulation in previous work (14). Second, a portion of the C-terminus (C1 region, amino acids 791-824) shown previously to be essential for SC assembly (14), is also critical for centromere coupling. Third, two mutants that are unable to assemble SC *(zip1-C2* and *zip1-M1*; (14)) are indistinguishable from wild-type cells for centromere coupling. We conclude that Zip1 contains some regions that are critical for centromere coupling but not SC formation and vice versa.

### The N-terminus of Zipl is essential for promoting the segregation of achiasmate partners

Though centromere coupling and centromere pairing both require Zip1, they have distinct genetic requirements suggesting they may operate by (at least partially) different mechanisms (20). To determine the regions of Zip1 that are required for achiasmate segregation we monitored the meiotic segregation of a pair of centromere plasmids that act as achiasmate partners in meiosis. Each plasmid carries an origin of DNA replication and the centromere of chromosome III, allowing the plasmids to behave as single copy mini-chromosomes in yeast. One plasmid is tagged with tdTomato-tetR hybrid proteins at a *tet* operon operator array (21), the other is tagged with GFP, as described above for chromosome I. Previous work has shown that such achiasmate model chromosomes disjoin properly in most meioses (22-24) and this segregation at anaphase I is correlated with the ability of their centromeres to pair late in prophase (5). To increase the synchrony of meiotic progression in this experiment *NDT80,* which promotes the transition out of prophase and into pro-metaphase, was placed under the control of an estradiol-inducible promotor (25-27). Meiotic cells were allowed to accumulate in pachytene of prophase, then induced to synchronously exit pachytene and enter pro-metaphase. We scored segregation of the plasmids in the first meiotic division by monitoring the location of their GFP and tdTomato tagged centromeres in anaphase I cells, identified by their two separated chromatin masses (Fig 3 A).

**Figure 3.**
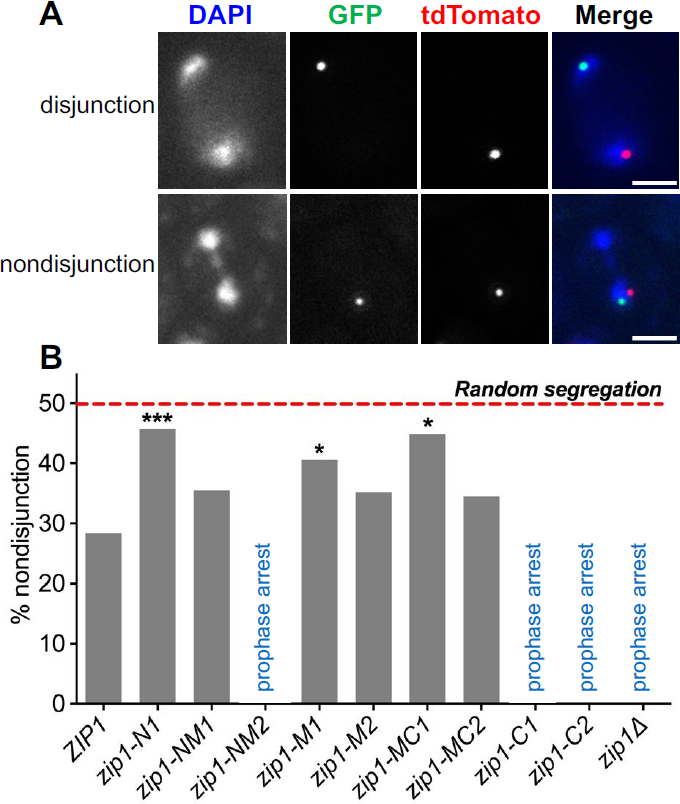
**Centromere plasmid disjunction requires the N-terminus of Zip1. A.** Representative binucleate cells with disjoined (a *ZIP1* cell) and non-disjoined (a *zip1-N1* cell) centromere plasmids. The segregation of CEN plasmids in anaphase I was assessed by monitoring the tetR-tdTomato and lacI-GFP foci localized to *tet* and *lac* operator repeats, respectively, inserted into a plasmid that contains 5.1 kb of *CEN3* sequence. **B.** Non-disjunction frequencies for CEN plasmids in each strain. n values: *ZIP1,* 250; *zip1-N1,* 190; *zip1 NM1,* 200; *zip1-M1,* 143; *zip1-M2,* 54; *zip1-MC1,* 69; *zip1-MC2,* 55. Statistical comparisons were performed with Fisher’s exact test to compare all genotypes to WT. Bonferroni’s correction was utilized to adjust for the number of comparisons. *p ≤0.05.; ***p ≤0.00625. Scale bars equal 2 µm.

Wild-type cells, under these conditions, exhibited 28% non-disjunction of the CEN plasmid pair (Fig. 3 B). The loss of Zip1 function can result in a pachytene arrest in some strain backgrounds (28) including the strain used in these experiments. Reducing the sporulation temperature to 23°C, as was done here, can permit a partial bypass of the arrest (28). Still several of the mutations (*zip1 Δ, zip1-C2, zip1-C1,* and *zip1-NM2)* yielded very few anaphase cells, and failed to sporulate, presumably due to the pachytene arrest. These observations are consistent with previously published work (14). Of the remaining mutants, the *zip1-N1* mutant showed significantly elevated non-disjunction of the centromere plasmids (Fig. 3 B). The *zip1-N1* mutant exhibits only mild defects in progression through meiosis, SC formation, sporulation efficiency, and the segregation of chiasmate chromosomes (14) and Figure S1), suggesting that amino acids 23-163 are more critical for mediating the segregation of achiasmate partners than for SC assembly and function.

Because achiasmate segregation is correlated with prior centromere pairing (7, 8), we tested whether the *zip1-N1* mutants were proficient in centromere pairing. Wild-type and *zip1-N1* cells containing the GFP and tdTomato tagged centromere plasmids were induced to sporulate and harvested five seven hours later when pachytene cells are prevalent. Chromosome spreads were then prepared and the distance between the tdTomato and GFP foci were measured in spreads exhibiting the condensed chromatin typical of pachytene cells (Fig. 4 A). The average centromere-centromere distance was significantly greater in *zip1-N1* mutants (Fig. 4 B) consistent with a loss of pairing. When spreads with an inter-centromere distance of less than 0.6 pm were scored as “paired” (see example in Fig. 4 A), the *zip1-N1* mutation was found to exhibit a significant reduction in the frequency centromere pairing between the achiasmate plasmids (Fig. 4 C).

**Figure 4.**
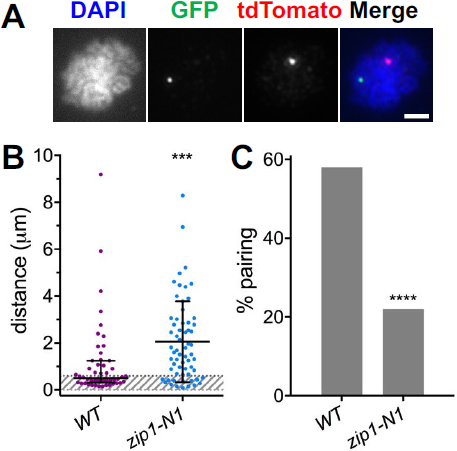
**Centromere plasmid pairing requires the N-terminus of Zip1.** Pairing of plasmid centromeres in prophase chromosome spreads was assessed by monitoring the pairing of tetR-tdTomato and lacI-GFP foci localized to *tet* operator and *lac* operator arrays on plasmids bearing a 5.1 kb region of chromosome *III*encompassing *CEN3.* **A.** An example of a spread with unpaired plasmid centromeres. **B.** Distances between the centers of the tdTomato and GFP foci in each spread (average and standard deviation). *** P=0.0002. The grey cross-hatched region indicates separation of less-than 0.6 µm between the centers of the foci, a distance used to infer pairing of the centromeres. **C.** The percent of spreads scored as “paired” in the *ZIP1* (58%, n=50) and *zip1-N1* (22%, n=63) strains. ****p<0.0001. Scale bar equals 2 µm.

### The N-terminus of Zipl is necessary for efficient localization to kinetochores

Failure of centromere pairing in the *zip1-N1* mutant could be due to a failure of Zip1 to associate with centromeres. To test this, we analyzed the co-localization of the Zip protein with kinetochores in *ZIP1* and *zip1-N1* strains. The experiments were done in a *zip4Δ* strain background to allow visualization of Zip1 localization independently of an SC structure. Images were collected using structured illumination microscopy and the level of co-localization was determined using ImageJ software (see Materials and Methods). Every *ZIP1* spread analyzed showed significantly more co-localization of Zip1 and Mtw1 than was found in a randomized sample (Fig. 5 A), consistent with earlier work (9, 10, 12), while many of the *zip1-N1* spreads showed no significant co-localization above the randomized control (Fig. 5 B). Consistent with these results, *zip1-N1* strains showed significantly lower levels of co-localization with Zip1 than was seen in *ZIP1* strains (Fig. 5 C).

**Figure 5.**
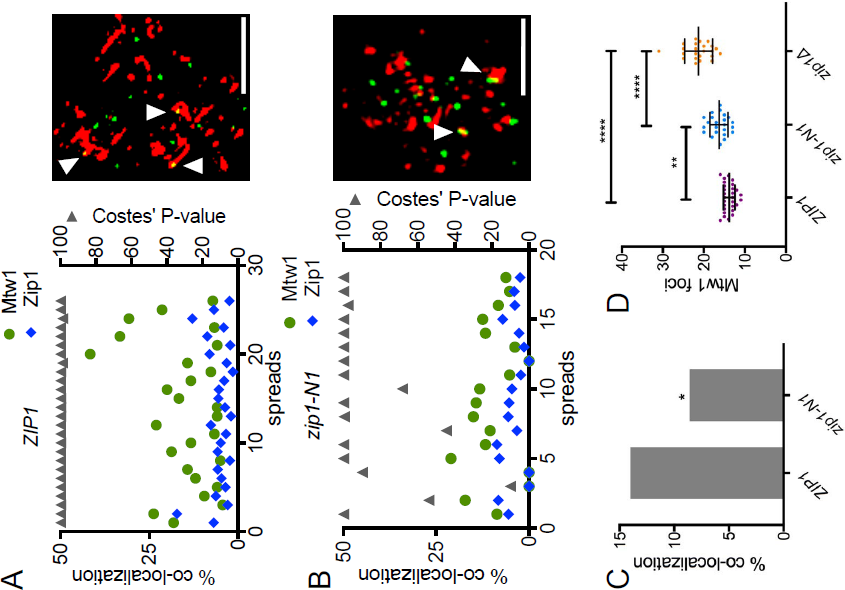
**The Zipl N-terminal domain is required for efficient co-localization to centromeres.** Chromosome spreads were prepared from prophase *ZIP1* and *zip1-N1* cells expressing Mtw1-GFP as a kinetochore marker. Indirect fluorescence structured illumination microscopy was used to visualize Mtw1-GFP and Zipl foci. **A and B.** The overlap of Mtwl foci with Zipl foci (green circles) and Zipl foci with Mtwl foci (blue circles) was measured in each spread and the statistical significance of the difference between the observed Mtwl co localization with Zipl from random simulations was evaluated with Costes’ P-value (gray triangles; greater than 95% is considered significant). Representative images from the two strains are shown. Zip1 (red), Mtw1-GFP (green), overlapping foci (white arrowhead), scale bars equal 2 µm. **C.** The average co-localization of Mtwl foci with Zipl across all the chromosome spreads was determined. * p<0.05. **D.** Centromere pairing was evaluated by counting the number of Mtwl-GFP foci in the chromosome spreads. *ZIP1* (n=27), *zip1-N1* (n=22), *zip1Δ* (n=22). **P<0.0l, ****P<0.000l.

### The N-terminus of Zip1 is necessary for the pairing of natural chromosomes

The reduced localization of Zip1-N1 protein to natural centromeres, above, and the failure of pairing of plasmid centromeres in *zip1-N1* strains (Fig. 4) raised the question of whether the *zip1-N1* mutation compromises the pairing of natural chromosomes. To assay centromere pairing we counted the numbers of kinetochore foci (Mtw1-GFP) in chromosome spreads from *ZIP1, zip1-N1* and *zip1Δ* cells, in the above experiment (Fig. 5) using structured illumination microscopy. Prior work had shown that in *zip4* mutants, with no SC, kinetochores are held in close proximity by centromere pairing. When *ZIP1* is deleted, the centromeres can resolve into two foci in chromosome spreads (29). The *ZIP1* strain gave an average of 13.9 kinetochore foci per spread, consistent with pairing of the 32 kinetochores. The *zip1-N1* mutant gave significantly higher numbers of kinetochore foci (average 16.4; p<0.01) signifying a loss of centromere pairing but not as dramatic a loss was observed in the *zip1* strain (average 21.3; p<0.0001).

## DISCUSSION

Our analysis of a set of in-frame Zip1 deletions has added to our understanding of the functional domains of the Zip1 protein, helping to ascribe particular Zip1 functions to specific regions of the protein. Zip1 is critical for SC assembly and processes that depend on SC assembly, including crossover formation and progression through pachytene (28). More recently it has become clear that Zip1 acts at centromeres both early in prophase, where centromeres become associated in a homology-independent fashion (centromere coupling), and later when homologous centromeres, or the centromeres of achiasmate chromosomes, become associated by remnants of the SC that remain at the centromeres after SC disassembly (reviewed in (30)). The experiments here were intended to clarify whether SC assembly, centromere coupling, and centromere pairing incorporate Zip1 in the same or different mechanisms, and if there are differences in the regions of Zip1 that are critical to each function.

### Centromere coupling and SC assembly

Prior work has shown convincingly that the structure that mediates centromere coupling is distinct from mature SC (9, 10, 20, 31). Several proteins (Zip2, Zip3, Zip4, Ecm11, Gmc2, and Red1) known to be essential for SC assembly are not required for centromere coupling. But the domains of Zip1 that are required for centromere coupling have not been defined. The experiments here reinforce that the requirements for Zip1 for centromere coupling and SC assembly are quite different. First, centromere coupling was proficient in *zip1-C2* mutants, which have severe defects in SC assembly. But these mutants exhibit little Zip1 expression, which may be due to the lack of a nuclear localization signal (32). Thus, this result is difficult to interpret other than to suggest that centromere coupling may require far less Zip1 than does SC assembly. Notably, the *zip1-M1* mutation, which also blocks SC assembly, is proficient in centromere coupling. The *zip1-M1* mutation, which eliminates amino acids 244-511, has a unique SC defect. The Zip1-M1 protein efficiently localizes to the axes of the homologous partners, but does not efficiently cross-bridge the axes (Fig. 1 C; (14)). This defect may reflect an inability of Zip1 molecules from opposite axes to associate with one another (as in Fig.1 B) or may reflect an inability of Zip1 to associate with central element proteins that promote or stabilize the cross bridging of axes by Zip1. In either case, such cross-bridging must not be important for centromere coupling, and is consistent with the finding that the central element proteins Ecm11 and Gmc2 are also not required for centromere coupling (31). Together these findings suggest that centromere coupling is probably not mediated by a structure that includes SC-like cross bridging. The only protein, beyond Zip1, that is known to be required for centromere coupling is the cohesin component Rec8 (9) (the requirements for the other cohesin subunits have yet to be reported). It may be that centromere coupling is mediated by the cohesin-dependent accumulation of Zip1 at early prophase centromeres (9, 29), followed by interactions between Zip1 molecules that promote the association of centromere pairs.

### Centromere pairing and SC assembly

Experiments performed mainly in a mouse spermatocyte model (3, 6) suggest that the SYCP1 (the functional homolog of Zip1) that persists at paired centromeres, after SC disassembly, is accompanied by other SC proteins. This suggests that centromere pairing could be mediated by a conventional SC structure. But the identity of regions of Zip1 that are critical for centromere pairing, and whether they are distinct from the regions necessary for SC assembly, have not been addressed. Our work suggests that there are significant differences in the requirements for Zip1 function in centromere pairing and SC assembly. We arrive at this conclusion following an evaluation of the centromere pairing phenotypes of the *zip1-N1* in-frame deletion. Prior work had shown this allele had no measurable differences from the wild-type *ZIP1* allele in spore viability, crossover frequency, and genetic interference, and a slight defect in the continuity of mature linear SC structures (14). In our strain background the *zip1-N1* mutation also exhibited wild-type levels of spore viability, and structured illumination microscopy confirmed the slight discontinuity in some SC structures in the *zip1-N1* background (Fig. S1). However, in centromere pairing assays the *zip1-N1* mutants showed major defects. In the *zip1-N1* mutant the centromeres of natural chromosome bivalents were more likely to become disengaged in chromosome spreads than was seen with wild-type controls, but the defect was not as severe as is seen in *zip1Δ* strains suggesting that there are regions outside of the N1 region that also promote association of the bivalent centromeres. It could be that these other regions are influencing things like cross-over frequency or distribution, that along with centromere-pairing help keep bivalent centromeres associated in the natural chromosome pairing assays. When we used achiasmate centromere plasmids, in which such functions cannot contribute to centromere association, then the *zip1-N1* phenotype becomes severe. The *zip1-N1* mutant showed a dramatic reduction in the pairing of plasmid centromeres. The fact that the Zip1-N1 protein is proficient for SC assembly but highly defective in centromere pairing suggests that the N-terminus imbues functions on the protein that are specifically required for centromere pairing. The mechanism of centromere pairing remains unclear as does the role of the Zip1 N-terminus, but kinetochore co localization experiments suggest that this region of Zip1 promotes localization to, or maintenance of, Zip1 at the centromeres in late prophase. The fact that early prophase centromere coupling is normal in *zip1-N1* mutants reinforces that coupling and pairing are fundamentally distinct processes and that the N1 region is not necessary for localization of Zip1 to centromeres in early prophase when coupling occurs.

### Meiotic prophase centromere pairing drives achiasmate disjunction

Experiments in yeast, *Drosophila* and mice have shown that SC-related proteins persist at paired centromeres after SC disassembly (3, 7, 8, 11). These observations have been the foundation for the model that centromere pairing promotes subsequent disjunction, especially of achiasmate chromosomes that are only connected at their centromeres. Demonstrating that this model is correct has been complicated by the fact that the SC is a central player in controlling meiotic progression. Thus, deletion of SC components, which eliminates centromere pairing, also impacts other processes such as synapsis, crossover formation, genetic interference, and the pachytene checkpoint, making it impossible to formally name centromere pairing, and not some other SC-related function as the driver of achiasmate segregation. The *zip1-N1* separation-of function allele, because it is largely wild-type for these other functions of Zip1, has made it possible to demonstrate in a compelling way that centromere-pairing in prophase is a requisite step in a process that mediates the segregation of achiasmate partners in anaphase.

The mechanistic question of how prophase centromere pairing drives disjunction remains to be answered. The fact that in yeast, mice and *Drosophila,* the majority of the centromeric SC components have been lost from the centromeres well before the partners begin to attach to microtubules makes this even more mysterious. The *zip1-N1* allele, which specifically targets centromere associations of Zip1, and the centromere pairing process, will be an important tool for addressing these questions.

## MATERIALS AND METHODS

### Strains

We created the same nine deletion mutants of *ZIP1* that Tung and Roeder had studied for their work in SC formation (14) by using standard PCR and two-step-gene-replacement methods (33, 34). All mutant versions of *ZIP1* were confirmed by PCR and sequencing. The native *ZIP1* promoter was unaltered in these strains allowing each mutant protein to be expressed at the appropriate level and time. Culturing of strains was as described previously (20). Strain genotypes are listed in Table S1.

### Centromere coupling assay

Centromere coupling was monitored largely as described previously (12). Cells were harvested five hours after shifting cultures to sporulation medium at 30°C. Meiotic nuclear spreads were prepared according to (35) with minor modifications. Cells were spheroplasted using 20 mg/ml zymolyase 100T for approximately 30 minutes. Spheroplasts were briefly suspended in MEM (100mM MES, 10mM EDTA, 500µM MgCl_2_) containing 1mM PMSF (phenylmethylsulfonyl fluoride), fixed with 4% paraformaldehyde plus 0.1% Tween20 and spread onto poly-L-lysine-coated slides (Fisherbrand Superfrost Plus). Slides were blocked with 4% non-fat dry milk in phosphate buffered saline for at least 30 minutes, and incubated overnight at 4°C with primary antibodies. Primary antibodies were mouse anti-Zip1 (used at 1:1000 dilution), rabbit anti-Zip1 (used at 1:1000 dilution; Santa Cruz y-300 SC-33733), rabbit anti MYC (1:400; Bethyl Laboratories A190-105A), mouse anti-MYC (used at 1:1000 dilution; gift from S. Rankin), chicken anti-GFP (used at 1:500 dilution; Millipore AB16901), rabbit anti DsRed (used at 1:1000-1:2000 dilution; Clontech 632496), and rabbit anti-RFP (1:500; Thermo Scientific 600-401-379). Secondary antibodies were obtained from Thermo Fisher: Alexa Fluor 488-conjugated goat anti-chicken IgG (used at 1:1200 dilution), Alexa Fluor 568-conjugated goat anti-mouse IgG (1:1000), Alexa Fluor 647 conjugated goat anti-rabbit IgG (used at 1:1200 dilution), and Alexa Fluor 568-conjugated goat anti-rabbit IgG (used at 1:1000 dilution).

Mtw1 (an inner kinetochore protein) foci (Mtw1-13xMYC) were quantified in spreads with an area of 15 µm^2^ or more to ensure centromeres were spread enough to assay. Centromere coupling would theoretically yield 16 kinetochore (Mtw1) foci while complete absence of coupling would yield 32 kinetochore foci. All strains were *spo11Δ/spo11Δ* to block progression beyond the coupling stage (12, 18). The individual performing the scoring was blinded to the identity of the mutation. The average number of Mtw1 foci seen in the chromosome spreads of each in-frame deletion strain was compared to the values obtained from the *ZIP1* and *ziplΔ* control strains, using the Kruskal-Wallis test, performed using Prism 6.0. The statistical data for the experiment are reported in Table S2.

### Achiasmate segregation assay

Non-disjunction frequencies of centromere plasmids were determined in a manner similar to previously published assays (7). Plasmids were constructed with arrays of 256 repeats of the *lac* operator or *tet* operator sequence inserted adjacent to a 5.1 kb interval from chromosome III that includes *CEN3.* These cells expressed a *GFP-lacI* hybrid gene under the control of a meiotic promoter and a *tetR-tdTomato* hybrid gene under the control of the *URA3* promoter. This produced fluorescent foci at the operator arrays (33, 34). Cells were sporulated at 23°C (rather than 30°C) as this has been shown to allow by-pass of the pachytene arrests triggered by some *ZIP1* mutations (28). Even at this temperature cells with the *zip1-C1, zip1-C2, zip1-NM2* and *ziplΔ* mutations mainly arrested in pachytene, so no anaphase segregation data were gathered for these strains. Harvested cells were either assayed fresh, or were frozen in 15% glycerol and 1% potassium acetate until the time at which they were assayed. Preparation for assaying the cells included staining the cells with DAPI and then mounting the cells on agarose pads for viewing as described previously (36). Anaphase I cells were identified by the presence of two DAPI masses on either side of elongated cells, indicating that the chromosomes had segregated. To avoid scoring cells with duplicated or lost CEN plasmids, only cells with one GFP focus and one tdTomato focus were assayed. Images were collected using the 100X objective lens of a Zeiss AxioImager microscope with band-pass emission filters, a Roper HQ2 CCD, and AxioVision software.

### Plasmid centromere pairing assay

Centromere pairing in pachytene was assessed using published methods (7) but with the centromere plasmids described above. Sporulation was done at 30°C. Chromosome spreads were prepared as described in (37), with the following modifications: Cells were harvested 5-7 hours after induction of sporulation at 30°C. After chromosome spreads were created and dried overnight, the slides were rinsed gently with 0.4% Photoflo (Kodak). Each slide was then incubated with PBS/4% milk at room temperature for 30 minutes in a wet chamber. Milk was drained off of the slide, and primary antibody diluted in PBS/4% milk was incubated on the slide overnight at 4°C. A control slide with PBS/4% milk was used for each experiment. The following day, the slides were washed in PBS, and incubated with secondary antibody diluted in PBS/4% milk for 2 hours in a wet chamber at room temperature. The slides were gently washed in PBS. DAPI (4',6-diamidino-2-phenylindole, used at 1µg/ml) was added to each slide and allowed to incubate at room temperature for 10 minutes. Slides were then washed gently in PBS and 0.4% Photoflo, then allowed to dry completely before a coverslip was mounted. Antibodies are described in the previous section. Only cells that exhibited “ropey” DAPI staining were scored in this assay, and were disqualified for assessment if there was more than one GFP focus or more than one tdTomato focus. In these cells, the distance between the center of the green focus and the center of the red focus was measured using AxioVision software. The distributions of distances in the *ZIP1* and *zip1-N1* strains were determined to be significantly different with the Kolmogorov-Smirnov test (Kolmogorov-Smirnov D=0.4032; P=0.0002) using the Prism 6.0 software package. As in previous work (7), foci with center-to-center distances less than or equal to 0.6 µm were scored as paired (these foci are typically touching or overlapping). The frequency of pairing (distance less than 0.6 µm) in the *ZIP1* (32 of 50) and *zip1-N1* (14 of 63) chromosome spreads was found to be significantly different (p<0.0001) using Fisher’s Exact test performed with the Prism 6.0 software package.

### Synaptonemal complex evaluation by structured illumination microscopy

Chromosome spreads were prepared according to the protocol of Grubb and colleagues (37) as described above, and harvested from sporulation cultures five hours after placing cells in sporulation medium at 30°C. To visualize the axial elements (Red1) and transverse elements (Zip1) of the SC by indirect fluorescence microscopy, chromosome spreads were stained with following primary and secondary antibodies: guinea pig anti-Red1 antibody (1:1000), goat anti-Guinea pig Alexa 488 antibody (Invitrogen) (1:1000), and rabbit anti-Zip1 antibody (1:800), donkey anti-rabbit Alexa 568 antibody (Invitrogen) (1:1000). Chromosome spreads were imaged with a Deltavision OMX-SR structured illumination microscope (SIM).

### Mtw1-Zip1 co-localization assay

Chromosome spreads were prepared according to the protocol of (37) as described above. All strains carried the *zip4Δ* to prevent SC assembly. Chromosomes were stained with primary antibodies: mouse anti-MYC (Mtw1-13xMYC) (Developmental Studies Hybridoma Bank) at 1:20 dilution and rabbit anti-Zip1 antibody at 1:1000 dilution and secondary antibodies Alexa 488 donkey anti-mouse (Invitrogen) at 1:1000 dilution and Alexa 568 goat anti-rabbit (Invitrogen) at 1:1000 dilution. Chromosome spreads were imaged with a Deltavision OMX-SR structured illumination microscope (SIM). Acquired images were converted to binary images using ImageJ software and the number of overlapping Mtw1-13xMYC and Zip1 foci were scored using the imageJ plugin, JACoP. To determine whether co-localization occurred at frequencies that were significantly higher than expected for random overlaps given the number of Mtw1 and Zip1 foci in each image, the foci in each image were randomized in one thousand simulations, then the frequency of random overlaps was determined and compared to the observed overlap. Costes’ P-value was then calculated to evaluate the statistical significance of the difference between the frequency of observed versus random overlap (38). In addition, the average co-localization observed for all of the *ZIP1* spreads (26 spreads, 238 Mtw1 foci, 33 co localized with Zip1) and all of the *zip1-N1* spreads (18 spreads, 279 Mtw1 foci, 12 co-localized with Zip1) was determined and the statistical significance of the difference determined using Fisher’s two-tailed exact test (p=0.0001). The experiment presented is one of two performed, both with the same outcome (significantly reduced Mtw1-Zip1 co-localization in the *zip1-N1* mutant).

### Centromere pairing of natural chromosomes

The chromosome spreads used in the experiment above were used to assay the number of distinct Mtw1-13xMYC foci in *ZIP1, zip1-N1* and *zip1Δ* chromosome spreads. With complete pairing of the homologous chromosomes, the thirty-two kinetochores should appear as sixteen Mtw1-13xMYC foci. In the absence of pairing, kinetochores from the paired homologs can sometimes separate far enough to be resolved as individual foci (the homologs remain tethered by crossovers and probably other constraints), thus giving higher numbers of Mtw1-13xMYC foci in theory up to thirty-two foci. The SIM images described in the preceding section were converted to binary images using ImageJ software and the number of Mtw1-13xMYC foci tallied for each spread using the Analyze Particles function in ImageJ. The average number of Mtw1-13xMYC foci per spread was determined for each genotype *(ZIP1, zip1-N1,* and *zipΔ1)* and the statistical significance of the observed differences between the genotypes was calculated with one-way ANOVA and multiplicity adjusted P values were obtained with Sidak’s multiple comparisons testing using Prism 7.0.

## Acknowledgements

We thank Marta Kasperzyk for the guinea pig Red1 antibody, Rebecca Boumil for the mouse anti-Zip1 antibody, and Jingrong Chen and Susannah Rankin for help with antibody preparation. We thank Lori Garman for help with statistical analysis. We are grateful to colleagues in the Program in Cell Cycle and Cancer Biology for helpful discussions of this work. The work was supported by NIGMS grant R01GM087377 to DD.

## Supporting Information Legende

**Figure S1. *zip1-N1* cells aeeemble eynaptonemal complexee and exhibit high epore viabilty.** Chromosome spreads were prepared from cells 5 hours after placing the cultures in sporulation medium and stained as described in Materials and Methods. The axial element protein is shown in green and Zip1 is shown in Red. Each panel presents representative spreads from A. ZIP1, B. zip1D and C. zip1-N1 strains. Panels to the right are larger images of individual chromosomes. The results in our strains are in keeping with the more comprehensive previous study of SC assembly in zip1-N1 mutants (Tung and Roeder, 1998) in that the zip1-N1 strain exhibited slightly less continuous Zip1 staining in pachytene-like spreads than was observed with the wild-type control strain. It is not clear if this reflects a slight reduction in assembly kinetics, or reduced continuity of the Zip1 in the mature SC of the zip1-N1 strain. D. Tetrads were dissected to assess spore viability in ZIP1 and zip1-N1 strains. Though in this sample set the zip1-N1 exhibited slightly lower spore viability than the wild-type control, as in prior studies (Tung and Roeder, 1998) there was no significant difference (Fisher’s exact test, P=0.83).

**Table S1. Strains Ueed in this Study**

**Table S2. Statietice for centromere coupling experimente**

